# Investigating the role of binding free energy, binding affinity and antibody escape in the evolution of SARS-CoV-2 spike protein

**DOI:** 10.1101/2022.10.15.512351

**Authors:** Matthew Young, Samantha J Lycett

**Affiliations:** University of Edinburgh

## Abstract

SARS-CoV-2 is considered a pandemic virus and presents a major strain on public health globally. SARS-CoV-2 infects mammalian cells by binding to its receptor, ACE2 which is mediated by the viral spike glycoprotein, specifically the receptor binding domain (RBD) within the spike protein. Recent development of vaccines against SARS-CoV-2 spike protein are currently the best strategy to reduce morbidity and mortality from infection. Like all viruses, SARS-CoV-2 evolves which may result in mutations which are benign or alter its viral fitness. The evolution of SARS-CoV-2 may increase the virulence, possibly by increasing the infectivity of the virus through strengthening the binding of the RBD to ACE2 or enabling the virus to evade naturally or vaccine induced immune responses. To address the need to characterise the evolution of SARS-CoV-2, this study has compared SARS-CoV2 sequences globally to the Wuhan reference strain at different time points. Additionally, by assigning scores to sequence data, which quantify each sequences binding strength to ACE2 and ability to evade patient derived antibodies, we have demonstrated that over time SARS-CoV-2 has evolved in less than one year to increase its ability to evade antibodies and increase the binding free energy between the RBD and ACE2.

## Introduction

Severe acute respiratory syndrome related coronavirus 2 (SARS-Cov-2) is a betacoronavirus that recently emerged within the human population in late 2019 causing over 150 million confirmed cases and over 3 million deaths to date (Gulyaeva and Gorbalenya, 2021). SARS-CoV-2 is related to other coronaviruses that infect humans such as SARSCoV and MERS-CoV, which both can cause high mortality (Lu et al., 2020). SARS-CoV-2 infection causes the acute respiratory illness, COVID-19 in most hosts, however, reports of chronically infected individuals have been recorded and the long-term impact of the infection has not been well studied (Johansson et al., 2021) (Choi et al., 2020). Like other coronaviruses, SARS-CoV-2 viruses are enveloped and contain a single strand positive sense RNA genome (Alexandersen, Chamings and Bhatta, 2020) that is relatively large spanning 29.8kb to 29.9kb in length (Khailany, Safdar and Ozaslan, 2020). SARS-CoV-2 genome encodes both non-structural (NSPs) and structural proteins. The NSPs perform important roles in the viral replication process, for example NSP12 is an RNA-dependent RNA-polymerase that ensures relatively accurate gene transcription (Chen, Liu and Guo, 2020). The four structural proteins encoded by SARS-CoV-2 are the membrane, nucleocapsid, envelope and spike proteins (Satarker and Nampoothiri, 2020).

SARS-CoV-2 mediates entry into human cells by binding to human Angiotensin convertase enzyme 2 (ACE2) and single-cell RNA-sequencing has identified that ACE2 is heavily expressed on type II alveolar cells explaining the viruses trophism for respiratory surfaces (Zou et al., 2020). SARS-CoV-2 binds to ACE2 by the viral spike glycoprotein, a homotrimeric protein and each monomer is comprised of S1 and S2 subunits (Kalathiya et al., 2020). The S1 subunit contains the amino acid (AA) residues that are in direct contact with human ACE2, known as the receptor binding domain (RBD) whereas the S2 subunit is required for membrane fusion between host cells and the virus (Huang et al., 2020). SARSCoV-2 is believed to have originated in an animal reservoir and was transmitted to humans in a spill-over event. The closest viral sequence matching SARS-CoV-2 is RaTG13, a coronavirus which has been sampled in bats with 96% sequence similarity however, the RaTG13 spike protein diverges from SARS-CoV-2 which suggests the presence of an intermediate host such as pangolins (Andersen et al., 2020), (Zhang, Wu and Zhang, 2020).

Viral evolution is stochastic and generates high levels of genetic diversity which may occur through several mechanisms such as mutations, recombination or reassortment resulting in viral variants (Arellano-Galindo et al., 2012). The evolution is however strongly influenced by selective pressures imposed by the environment, which select for variants with the optimal viral fitness. Within a host, the evolution of a virus is often determined by its ability to evade the immune response that recognises specific viral epitopes (Boni, Gog, Andreasen and Feldman, 2006). Viruses, particularly RNA viruses replicate with a relatively high mutational burden (Peck and Lauring, 2018), which often result in progeny that vary slightly antigenically. These mutations may enable specific viral variants to evade the host immune responses and therefore have a competitive advantage over other variants. Over time the antigenic landscape of viruses can change because of immune evasion, this is known as antigenic drift (Earn, Dushoff and Levin, 2002). Viral evolution is of particular concern as treatments such as vaccines and therapeutic drugs are developed to act against specific viral targets and viral evolution can therefore decrease their efficacy. Antigenic drift occurs in Influenza A virus (IAV), which has a high evolutionary rate and due to this evolution, IAV H3N2 vaccines are required to be updated annually to remain an effective control measure. Additionally, a single nucleotide polymorphism (SNP) in H3N2 can result in resistance to the antiviral adamantane (Nelson et al., 2009), which demonstrates the importance in surveying the prevalence of mutations. Mutations can increase the virulence of a virus, therefore increasing the potential of a virus to increase mortality or infectivity (Long et al 2020). demonstrating the importance in surveying the prevalence of mutations. The mutation rate of SARS-CoV-2 is relatively low for RNA viruses as determined by a molecular clock (Duchene et al., 2020), however many variants have been reported and it is not yet known if many of these mutations have significant impacts on viral behaviours and if the newly developed SARS-CoV-2 vaccines will need to be updated to remain effective.

Recently, several mutations have arisen in SARS-COV-2, some of which are considered variants of concern, as they are believed to have an increased potential for infectivity (Zhang et al., 2020) (Faria et al., 2021) many of these mutations are found within lineages such as P1 or B.1.351 (PANGO nomenclature) (Rambaut et al, 2020). Increased infectivity can be the result of many possible mechanisms, however the aetiology of mutations within the spike protein specifically that increase infectivity are likely due to an increased capacity of the RBD to bind to ACE2 and an increased potential to evade the host immune response. Therefore, there is a need to assess the ability of RBD mutants to bind to ACE2 and evade patient derived antibodies to determine how infectious specific variants are and if SARS-CoV-2 is evolving to become more infectious within humans. The first aim of the study was to investigate the level of genetic diversity globally at different time points and then examine the specific mutations occurring in different nations over time. Following this, viral sequences will be assigned scores that represent their individual ability to evade antibodies, and capacity to bind to ACE2 to measure the relative infectivity of SARS-CoV-2 over time. Using this data, the evolution of SARS-CoV-2 can be better understood.

## Methods

### SARS-CoV-2 sequence data

SARS-CoV-2 spike sequence data was downloaded from the GISAID database. From the dataset, sequences from the sample nations Australia, Brazil, Japan, South Africa, Spain, United Kingdom, and the USA were extracted and arranged into subsets based on the date of sequencing. These countries were selected as they had the greatest viral sequencing efforts in each continent (excluding Antarctica) (Furuse, 2021). Subsets of data from each country were arranged into three different waves of infection: Wave 1 included sequences from March, April, and May 2020; Wave 2 included sequences from August, September and October 2020, and Wave 3 included sequences from December, January, and February 2020-2021. Any sequences containing undefined AAs denoted by the presence of “X”‘s in residues 331-526 were removed from analysis. Sequences containing INDELS were realigned with the Wuhan reference sequence (Genbank accession: NC_045512) and reincorporated into the analysis. All sequence data were aligned using the MUSCLE algorithm using the software MEGA. In total, 85,666 complete SARS-Cov-2 spike protein sequences were suitable for this study.

### Entropy calculations

Shannon information entropy can be used to measure the degree of diversity within a system (Schneider, Stormo, Gold and Ehrenfeucht, 1986). Shannon entropy can quantify the diversity in AA composition at sites from sequence data. In this study, Shannon entropy (H) at an AA site position (X) is calculated as a function of the frequency of the different types of AA:

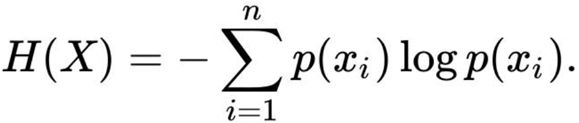

In which *i*=1-20 is the AA identity at each position. Shannon information entropy was derived from sequence data gathered from the sample nations at waves 1, 2 and 3. The calculation was applied to all sequence data within the software “R”. The average entropy per position in the sample nations at waves 1, 2 and 3 were then calculated to gather the average entropy of sequence data in different waves. AA sites with a high entropy value indicate heterogeneity in AA compositions between sequences at this site and indicate diversification. Whereas a low entropy value of 0 indicates a homogenous AA composition at a given site which may indicate if an AA is fixed within a population.

### Mutation screening

To identify the specific sites of mutations within each individual sample nation at different time points, the sequence data was compared to the Wuhan reference strain. Spike protein residues in which the AA composition differs from the Wuhan reference strain were plotted, residues in which the AA composition differed from the Wuhan reference strain in more than 10% of the sequences sampled were highlighted and included in table 1. Analysis of AA composition at certain sites was determined by plotting the frequency of AA composition over time using the software “R”

**Table 1.**
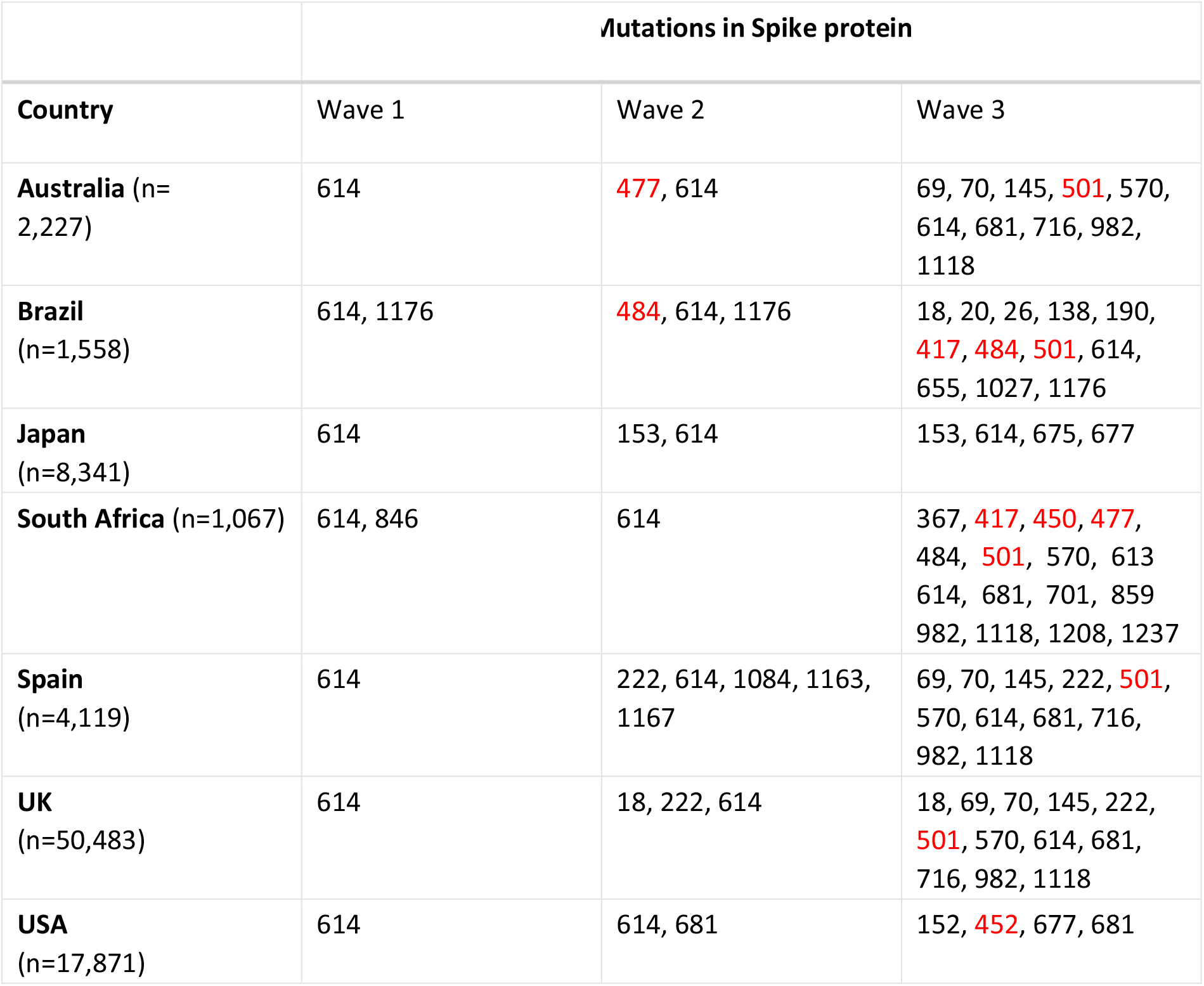
Positions in the spike protein where mutations from the Wuhan reference sequence are found in more than 10% of the sequences from each sampled nation in waves 1-3. Amino acid residues coloured in red denote those found within the RBD of the spike protein. The total number of sequences from each country in waves 1-3 are listed under the country names.

| **Table 1** | **Mutations in Spike protein** |  |  |
| --- | --- | --- | --- |
| **Country** | Wave 1 | Wave 2 | Wave 3 |
| **Australia** (n=2,227) | 614 | **477**, 614 | 69, 70, 145, **501**, 570, 614, 681, 716, 982, 1118 |
| **Brazil** (n=1,558) | 614, 1176 | **484**, 614, 1176 | 18, 20, 26, 138, 190, **417**, **484**, **501**, 614, 655, 1027, 1176 |
| **Japan** (n=8,341) | 614 | 153, 614 | 153, 614, 675, 677 |
| **South Africa** (n=1,067) | 614, 846 | 614 | 367, **417**, **450**, **477**, 484, **501**, 570, 613, 614, 681, 701, 859, 982, 1118, 1208, 1237 |
| **Spain** (n=4,119) | 614 | 222, 614, 1084, 1163, 1167 | 69, 70, 145, 222, **501**, 570, 614, 681, 716, 982, 1118 |
| **UK** (n=50,483) | 614 | 18, 222, 614 | 18, 69, 70, 145, 222, **501**, 570, 614, 681, 716, 982, 1118 |
| **USA** (n=17,871) | 614 | 614, 681 | 152, **452**, 677, 681 |

### Scoring Sequences

Sequences were assigned individual scores that quantify the binding free energy (BFE) of the RBD to ACE2, the binding affinity of the RBD to ACE2 and the ability of mutations at each site to evade antibodies (TSE score) (see next section). Sequence data were compared to the Wuhan reference strain and every AA in the test sequences that differed from the reference sequence between positions 333 and 526 were allocated these scores according to the position of the AA mutation. Where multiple mutations were present in a sequence, the scores were averaged across the sequence. Scores were derived from multiple datasets that were modified for this study.

### TSE scores

Greaney et al, (2021) measured the ability of mutations within the RBD to escape patient derived antibodies. All possible AA mutations in the receptor binding domain were mapped onto a yeast surface display platform and were then titred against sera derived from 11 patients infected with SARS-CoV-2 approximately 114 days post symptom onset. The heterogeneity in antibody binding to SARS-CoV-2 that is reflected by variable total antibody escape scores between subjects 1 to 11 justifies the use of this data as a measure of a mutations ability to evade antibodies within a host population. The average total antibody escape score per site (TSE) was then calculated between all patients to produce a model of population level immunity for use in this study. Antibody escape scores were missing for several positions within the RBD (n=24) and therefore antibody escape scores could only be assigned to mutations in sequences where a score was available for that particular position.

### Binding affinity and BFE scores

Binding affinity data was calculated by Starr et al (2020) who mapped every possible AA mutation within the RBD onto a yeast surface display platform. The binding affinity was calculated by measuring the dissociation constant of each RBD variant to ACE2. The binding affinity dataset was selected as it could provide a measure of infectivity through quantifying the binding affinity of RBD variants to ACE2. This study adapted this data for scoring viral sequences by calculating the average change to binding affinity induced by all mutations at each site within RBD positions 333-526. BFE data was calculated by Chen et al (2021) who integrated topological representation and, machine learning to predict the BFE between protein-to-protein interactions. Chen et al (2021) calculated the BFE between SARS-CoV-2 RBD and ACE2 for every possible AA substitution at residues 333-526. The BFE dataset was selected as the BFE between RBD and ACE2 was proportional to infectivity of SARSCoV (Forouzesh and Mishra, 2021) and their approach was found to be 22% more accurate at predicting BFE from protein-to-protein interactions than previous datasets (Wang, Cang and Wei, 2020) and therefore was suitable for use in this study. This study calculated the average change to BFE that mutations cause at residues 333-526, this produced a BFE score for each position.

## Results

### Entropy

To examine the increase in diversity of the spike protein over time, the Shannon entropy was calculated across all the samples for all seven country data sets for waves 1, 2 and 3 respectively (figure 1). This indicates that the Shannon entropy increased over the successive waves, meaning that the spike sequence diversity increased over time. Across the three waves, position 1274 appears to have consistently high entropy values, however this is due to poor sequencing quality at terminal ends of genome and therefore will be excluded from analysis. In wave 1, there are two positions with high entropy scores, 614 and 1176. At wave 1, position 614 has an entropy value of 0.4032, indicating that there is heterogeneity in the AA composition at this site. In wave 1, the entropy of positions located within the RBD (333-526) did not rise above 0.05 and therefore there were few AA substitutions occurring within the RBD during this time.

**Figure 1.**
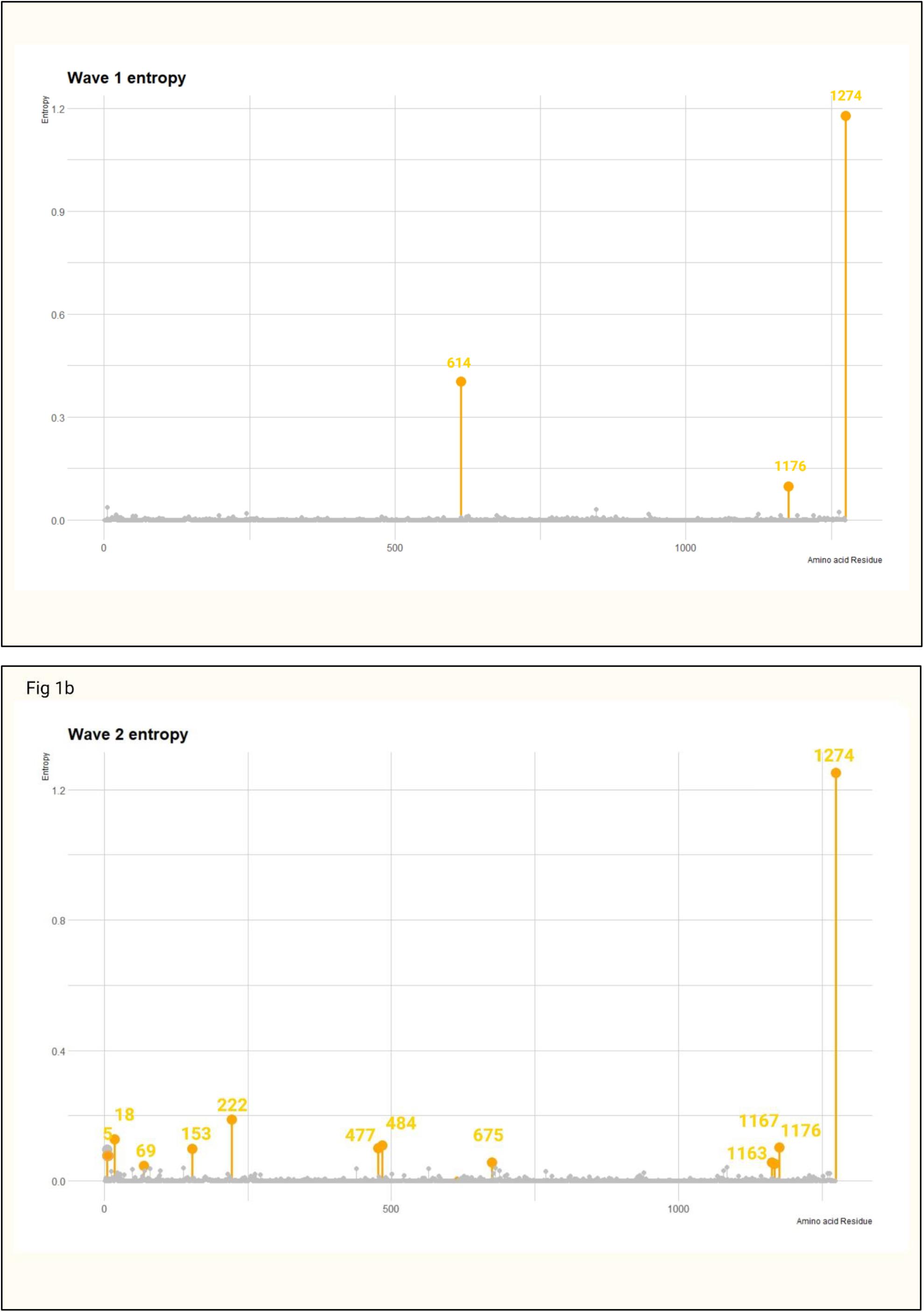
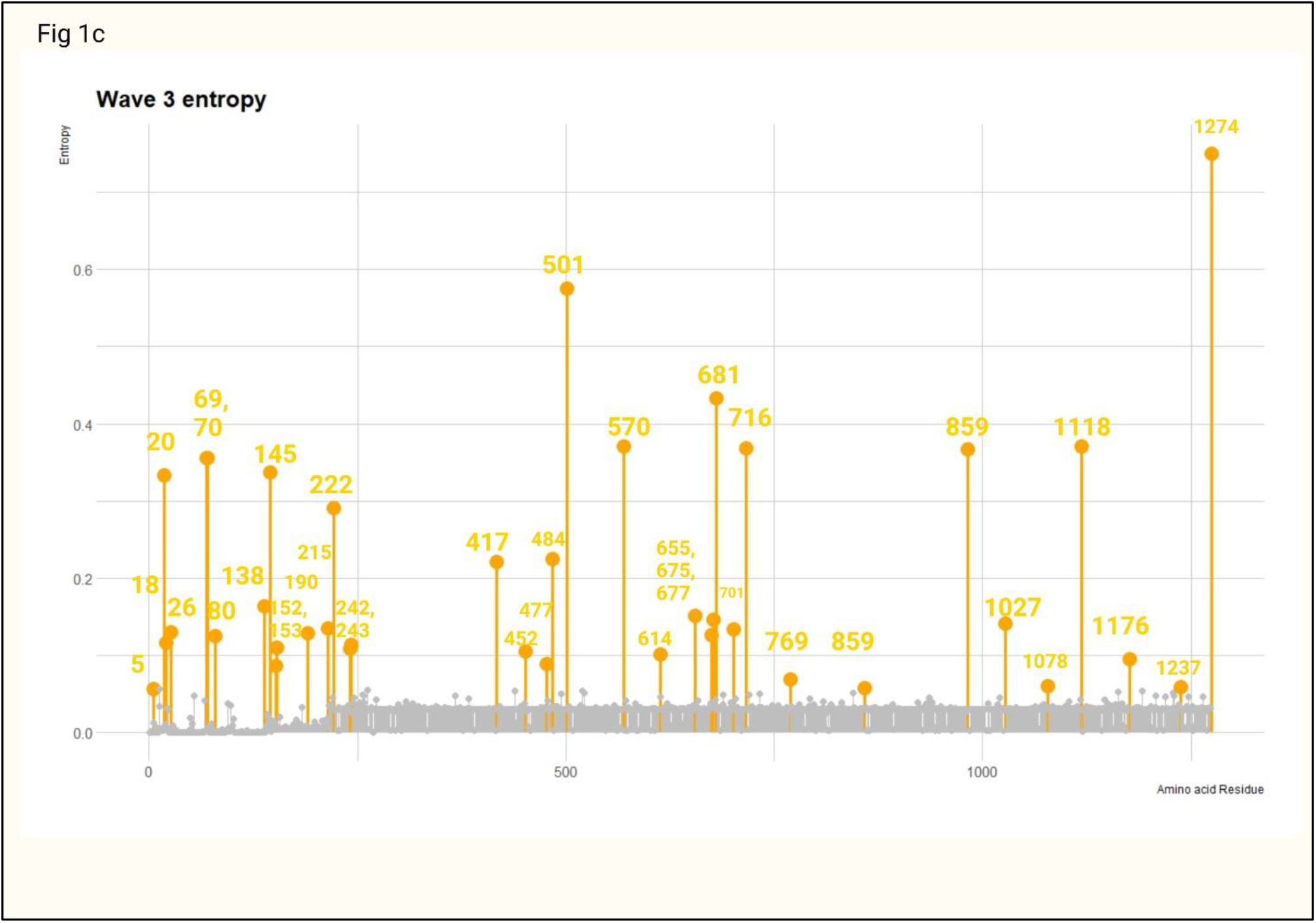
Entropy of SARS-Cov-2 in sample nations. Fig1**a**, average entropy across all sampled nations in wave 1. Fig 1**b**, average entropy across all sampled nations in wave 2. Fig1**c** Average entropy across all sampled nations in wave 3.

In wave 2, the entropy values have increased in a number of positions within the spike protein in comparison to wave 1 with positions 222, 477 and 484 having the greatest entropy values during this time. The entropy at both positions 69 and 70 has increased during wave 2 with values of 0.046 and 0.032 respectively. The entropy value at position 614 has decreased from 0.4302 during wave 1 to 0.0063 in wave 2, this suggests that a mutation occurred at some stage before or during wave 1 and subsequently became fixed in the population by wave 2.

By wave 3, more positions within the spike protein have increasing entropy values with 8 positions within the RBD (333-526) having entropy greater than 0.05, which suggests that over time spike protein is evolving. The entropy value at position 501 has sharply increased from 0.0218 in wave 2 to 0.5759 in wave 3, which may imply certain AA substitutions at this position enhance viral fitness or infectivity. In addition, the entropy values at positions 69 and 70 have increased to 0.3558 and 0.3553 respectively, the close proximity of both these positions and entropy values supports the conclusion that a mutation is equally affecting the AA composition in both positions at the same time. The entropy value at position 1176 in wave 3 was 0.0954, the entropy at position 1176 has remained consistent between waves 1 and 3 and therefore it may be deduced that this is because mutations at 1176 are consistently appearing or perhaps the mutation at this site does alter viral fitness significantly and the spreading is the result of a founder effect. Calculating the entropy of the sequence data has demonstrated positions in which the AA composition are diversifying, however in order to identify which mutations in the spike protein have occurred in each country during each wave, the sequence data from each nation needs to be compared to the Wuhan reference genome.

### Mutation analysis

By analysing sequence data from seven nations individually, at three different waves of infection, analysis can reveal all mutations within the spike protein and the AA compositions at each site when compared to the Wuhan reference genome. In figure 2, the spike protein residues where the AA composition differs from the Wuhan references genome in more than 10% of samples are highlighted in red. Figure 2 shows the results for Brazil, please see the appendix for the results from the other countries.

**Figure 2.**
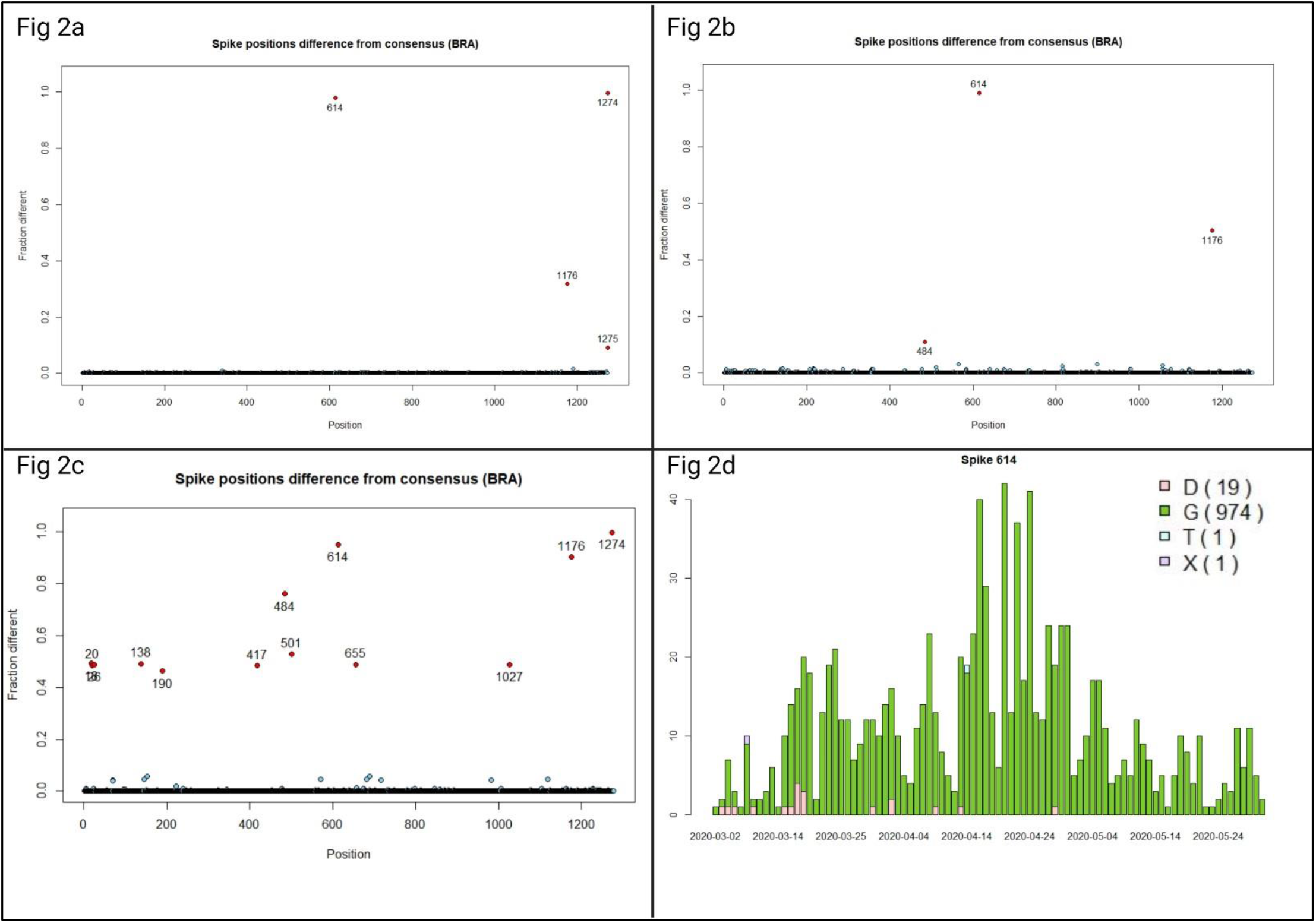
Amino acid positions which differ from the Wuhan reference sequence in each wave and the amino acid composition at site 614. Figure 2**a**, 2**b** and 2**c**, depict the sequences from wave 1, 2 and 3 respectively and amino acid positions in which the composition differs from the Wuhan reference sequence in more than 10% of sequences sampled are highlighted in red and labelled. Fig 2d. Barplot of the amino acid composition at position 614 in brazil during wave 1.

Between March and May 2020 (Wave 1), the results of figure 2a demonstrate that in Brazil, the AA found at residue 614 differed from the AA composition in the Wuhan reference strain in nearly 100% of the samples sequenced at this time. Further analysis of site 614 in figure 2d demonstrates the heterogeneity in AA composition in early March 2020. An AA substitution from aspartic acid (D) at position 614 in the reference strain to glycine (G) (denoted D614G) is observed early in brazil wave 1, which is the result of a SNP. Figure 2 shows that over time this AA substitution at position 614 has become fixed. Mutations prevalent in other nations are depicted in table 1. In the six other nations analysed (see appendix), the AA identity at position 614 of the spike protein differed from the Wuhan reference in approximately 40% (Australia) to 90% (south Africa) of the sequences sampled in wave 1. The varying prevalence of the mutation at position 614 at wave 1 is also demonstrated by figure 1(a). By wave 2, the AA composition at position 614 differed to the reference sequence in 100% of all sequences sampled in all seven nations examined, indicating that the D614G mutation was the first mutation within the spike protein to occur in SARS-CoV-2 within the human host and become fixed within the viral population.

In Brazil at wave 2, figure 2b reveals that mutations are occurring at position 484 in more than 10% of the sequences sampled. Analysis of the AA composition at this time point reveals glutamic acid (E) and lysine (K) are found at this site. Mutations at position 484 are not observed in any other nations at this time point, indicating that the mutation originally occurred in Brazil.

The prevalence of the E484K mutation in Brazil in wave 3 has increased to approximately 20%, the growing prevalence of this mutation along with the position within the RBD suggests that this substitution may enhance viral fitness within the population. An additional mutation within the RBD occurred in wave 2, as the S477N substitution was dominant in over 90% of sequences sampled from Australia, however this mutation was not seen in Australia in wave 3. The low frequency of the S477N mutation in wave 3 in Australia, may imply that the mutation causes a substantial cost to viral fitness however it is more likely that strenuous lockdown restrictions imposed within Australia may have limited the opportunity for transmission of viruses containing this variant, which are reflected by the low availability of sequences at this time (n=188). Therefore, this analysis highlights complexities of mutations within the RBD warrants further investigation into the infectivity of mutations within the RBD.

The nucleotide mutation at position 1176 causing the substitution V1176F have been consistently observed within Brazil in waves 1-3. At wave 2, AA compositions of Valine and phenylalanine were approximately homogenous at position 1176 however by wave 3, 1176F dominated. Residue 1167 is located in the S2 subunit and therefore mutations at this site may alter membrane binding between the virus and host cell which may alter infectivity. Despite its potential for increased pathogen fitness and several independent occurrences, this mutation has not been seen in more than 10% of sequences sampled from any other country at any other time point.

Table 1 indicates that a mutation has occurred in both positions 69 and 70 during wave 3 in Australia, Spain, and the UK. Analysis of the AA composition at these sites reveals a mutation causing an AA deletion has occurred, the approximately equal entropy values in figure 1 supports the deduction that deletions at these sites are associated with each other. The presence of this mutation in several countries further suggests that positions 69 and 70 are recurrent deletion regions of the spike protein. In wave 3, there was a significant increase in the number of AA mutations which have occurred in the spike protein in all nations. While the only mutation to become fixed within all sampled populations remains the D614G mutation, the fraction of sequences which contain other mutation increases, suggesting that they may become fixed in the future. By wave 3, in all countries except for Japan, several mutations have occurred within the receptor binding domain. In South Africa, there are six mutations prevalent in circulating viruses that lie within the RBD, which is of particular concern and adds to the need to greater analyse the effect of mutations within the RBD. Therefore this analysis has highlighted that understanding the effect of mutations which arise within the RBD are key in to understanding the infectivity of viral variants and may reveal evolutionary patterns of this virus.

### Sequence scores

Sequences from each nation were assigned scores to quantify their ability to bind ACE2 and evade antibodies. This approach has revealed patterns in the evolution of SARS-COV-2 in nearly one years’ worth of sequences. In all nations sampled, the average BFE score of each sequence increased between waves 1 and 3. The increase in BFE from sequences in wave 1 to wave 2 were modest and this likely due the presence of only few mutations within the RBD within this narrow time frame which is consistent with the relatively low mutation rate of SARS-CoV-2. The mutations that did arise in sequences in wave 2 that increase BFE between the RBD and ACE2 were likely selected for as a result of their increased infectivity. Sequences from wave 3 have the greatest average BFE score compared to other time points, which demonstrates that increased BFE between RBD and ACE2 may enhance viral fitness or infectivity. Therefore, over time sequences from all countries are acquiring mutations within the RBD which increase the BFE of SARS-COV-2 to ACE2. This results in sequences that have an increased potential for infectivity. However, the binding affinity scores derived from Starr et al (2020) suggest that the mutations within the RBD between waves 1-3 on average do not strongly affect the binding affinity of the RBD to ACE2 and may even decrease binding affinity.

Additionally, figure 3 shows that between waves 1 and 3, the average total antibody escape score at each RBD site (TSE) of SARS-CoV-2 increases. In all nations at wave 1, the average score representing each sequence’s ability to evade antibody binding is close to zero, indicating that selective pressures imposed by the hosts antibody response did not impede viral transmission as to be expected from a novel virus. By wave 2, the ability of SARS-CoV-2 to escape antibody binding has increased as seen by a modest increase in the average TSE score in sequences at this time, which is likely due to increasing seroprevalence against SARS-CoV-2. By wave 2 a large proportion of populations remained immunologically naive to SARS-Cov-2, possibly due to extensive lockdown restrictions, therefore the evolutionary pressures selecting for mutations with high TSE scores, are not as high as they were in wave 3. Brazil and South Africa both contain sequences in wave 3 that contain the greatest average TSE score, whereas sequences from Spain and the USA do not have as high average TSE scores as other nations. With greater evolutionary pressures to evade antibody mediate immunity in wave 3, lineages such as P.1 in Brazil that contain variants conferring higher levels of antibody escape are selected for over lineages such as A.2.5 which do not contain as many mutations from the Wuhan reference sequence. This analysis has revealed that SARS-CoV-2 lineages present globally in wave 3, can evade antibody binding from sera derived from SARS-CoV-2 infected patients; therefore, suggesting antigenic drift of spike protein epitopes which may have implications on vaccine efficacy. Overall, these results have shown that SARS-CoV-2 is evolving to increase BFE between ACE2 and RBD and increasing its ability to evade antibodies.

**Figure 3.**
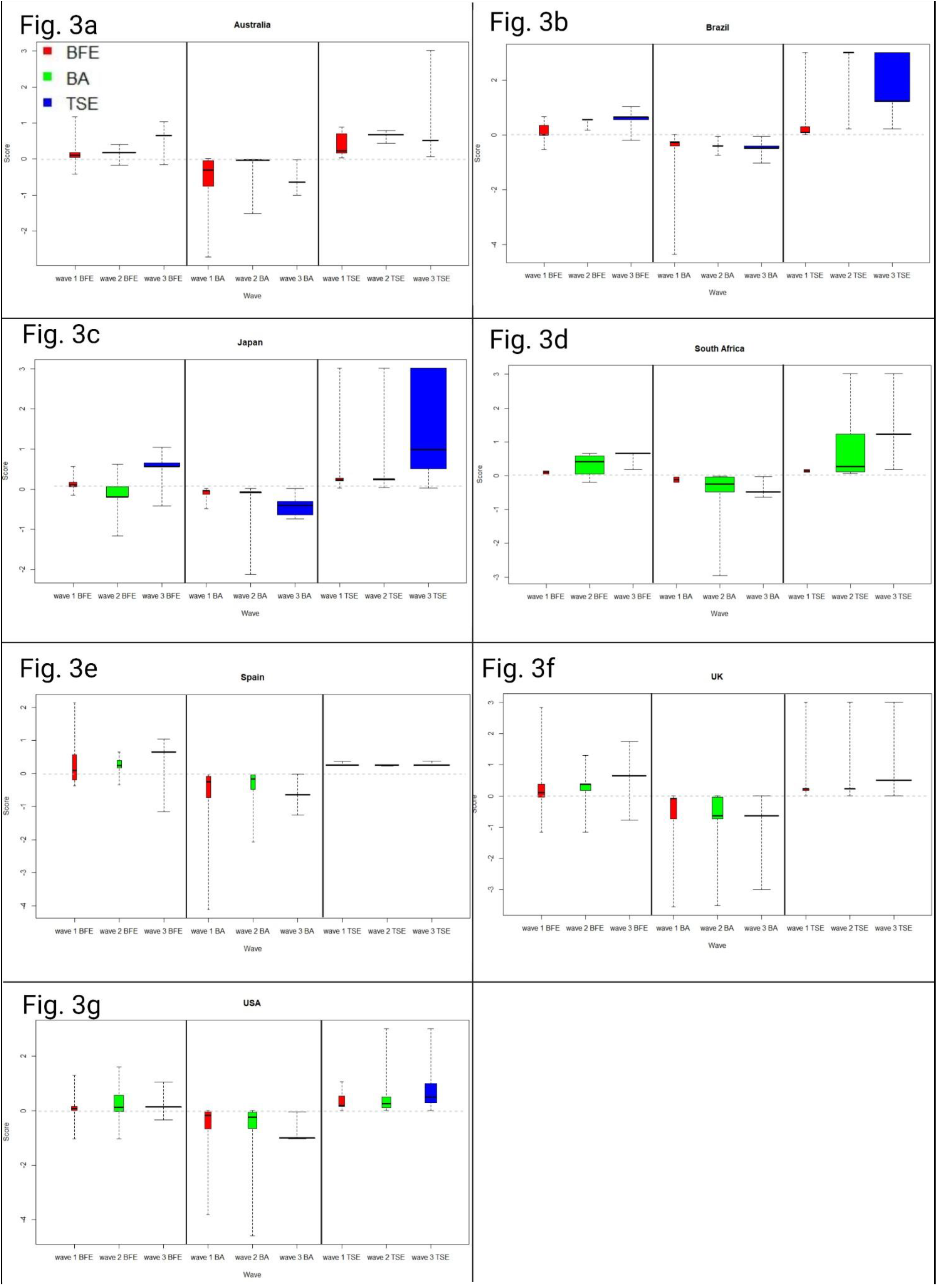
Boxplots of the average score of each mutation from sequences in sample nations between waves 1-3. The boxplots coloured red are the binding free energy between ACE2 and RBD scores (BFE), the green boxplots are the binding affinity between ACE2 and RBD scores (BA) and blue boxplots are total site escape of mutations from antibodies scores (TSE).

## Discussion

This study has analysed the sequence data of SARS-CoV-2 to determine the prevalence of mutations within the viral spike protein using entropy methods and direct comparison to the Wuhan reference sequence. The evolution of SARS-CoV-2 RBD has been characterised by developing a scoring system which may suggest increased infectivity of the virus. Calculating the Shannon entropy of spike protein residues of SARS-CoV-2 has revealed sequence diversity increases over time. Figure 1 demonstrates that between waves 1-3 the positions with the greatest entropy; therefore, the greatest diversity are clustered in three discrete regions of the spike protein. These regions are between approximately residues 5190, 390-680, and 980-1240. Residues 5-190 are found within the N-terminal domain in the S1 subunit and are critical in host-receptor binding (Saputri et al., 2020). Additionally, residues 390-680 span a proportion of the RBD and high diversity within these regions is of particular concern as the increasing BFE scores suggest that overall, the spike protein is evolving to strengthen binding to ACE2. Diversity among positions within these discrete regions within the spike protein that is observed from entropy values is also seen in Brazil by wave 3 (figure 2c). Direct comparison of sequence data from each nation at specific samples provided greater resolution into what specific mutations occurred in each nation at what time. In this study the E484K substitution first appeared within sequences from Brazil in wave 2 where it is believed it arose independently from where the mutation was first detected in the South African lineage B.1.351 (Hodcroft et al, 2021). E484K was not prevalent enough in wave 2 sequences within South Africa to be time to be detected by this analysis. Shannon entropy at position 1176 remained consistent, with values at 0.097, 0.103 and 0.0954 in waves 1-3 respectively and the AA composition at position 1176 indicates the V1176F substitution did not cause an apparent selective advantage that can be implied through measuring the frequency. Studies have suggested that the V1176F is strongly associated with increased mortality (Lange et al., 2020), however these studies are subject to sampling bias. This highlights the need for further characterisation of mutations such as V1176F. By quantifying SARS-CoV-2 evolution by scoring viral sequences we have demonstrated that SARS-CoV-2 has evolved to increase the BFE of the RBD to ACE2. Chen et al (2020) has described how the BFE between ligand and receptors is proportional to infectivity, therefore this approach has shown that SARS-CoV-2 evolves to become more infectious. Current trends of SARS-CoV-2 RBD evolution suggests that mutations conferring greater BFE between the RBD and ACE2 are likely to occur in the future and will be more infectious than other strains, highlighting the need to enhance genomic surveillance particularly for SNPs resulting in AA substitutions at these sites. AA substitutions at positions 489, 452, 422, 342 and 444 result in the greatest increase to BFE between the RBD and ACE2, and therefore are of particular interest. The L452R substitution has already been observed from the analysis of mutations within sequences from the USA in wave 3 as seen in table 1. The mutation is only prevalent in approximately 5% of sequences in the USA in wave 3 however this will likely increase in the future.

This approach has also revealed that the ability of SARS-CoV-2 lineages from all sample nations to evade antibodies has increased in only one year. Scoring sequences abilities to evade antibodies has identified variants that can evade antibodies and therefore pose a threat of diminishing the efficacy of vaccine induced immunity. Sequences from South Africa in wave 3 demonstrated the greatest ability to evade antibodies as seen by the large TSE scores at this time. During wave 3 in South Africa several lineages were circulating, these are the B.1.1.7 lineage containing the RBD mutation N501Y, the B.1.351 lineage containing the RBD mutations N501Y, E484K and the P.1 lineage containing the RBD mutations K417T, E484K and N501Y. Viral neutralisation assays of B.1.351 variants reported a 7.6 fold and 9 fold reduction in antibody neutralisation from patients inoculated with the PfizerBioNTech and Oxford-AstraZeneca vaccines respectively compared to an early Wuhan isolate (Zhou et al., 2021). Therefore, it may be cautiously inferred that a TSE score greater than 1 may cause a 7.6- and 9-fold reduction in the neutralisation ability of Pfizer-BioNTech and Oxford-AstraZeneca vaccines. In addition, the low occurrence of mutations occurring in Japan and the USA as seen in table 1 may suggest that viral transmission in these nations has been facilitated without strong selective pressures compared to other nations which may in fact be demonstrated by their relatively low increases in antibody escape scores in waves 1-3. Additionally, observations of single sites in which mutations cause high antibody escape support claims of B-cell immunodominant sites within the spike protein (Zhang et al., 2020). Currently the cut-off vaccine induced antibody neutralisation titre that maintains protection against SARS-CoV-2 is unknown and will be required to determine at which point vaccines are required to be updated. Vaccines inoculating against IAV subtype H3N2 are updated annually, due to antigenic drift of the haemagglutinin protein which can result in approximately 3 mutations becoming fixed per year (Fitch, Bush, Bender and Cox, 1997). Currently, the only mutation within the spike protein that has become fixed within the population is the D614G substitution suggesting the SARS-CoV-2 vaccines do not need to be updated annually, however 37 other mutations have been identified within this study (table 1), and the increasing prevalence suggests that several of these mutations such as N501Y, E484K and L452R may become fixed within a year.

## Limitations

To score the ability of mutations to escape antibodies, a model of population immunity was produced from data that measured the ability of mutations at particular sites to escape binding from sera derived from 11 patients infected with SARS-CoV-2. However, this data does not consider the effect of cytotoxic T-cells or other immune effector responses involved in viral recognition or clearance. Additionally, missing data for the effect of mutations in specific sites (n=24) meant that certain mutations could not be assigned a score and likely led to an underrepresentation of the true increase in TSE over time. Additionally, the sequencing capacity globally varies with South African analysis containing only 1,607 sequences overall whereas there were 50,483 sequences derived from the UK alone. However, as all sequence data were analysed relative to their respective nation, this does not have a significant impact on the overall analysis. The scores for BFE and binding affinity of the RBD to ACE2 may not be an accurately represent *in-vivo* conditions as the BFE scores were calculated *in-silico* and the binding affinity scores were calculated from an *invitro* experiment. Furthermore, all scores assigned to sequences were the average value of each individual score for specific AA substitutions at that site and therefore scores assigned to each mutation are an average of all possible scores at that site rather than the specific mutation.

### Future Directions

This analysis has presented an opportunity to score SARS-CoV-2 sequences based on their potential for infectivity. Future study with this approach should be integrated into growth rate models to develop a greater appreciation of the infectivity of SARS-CoV-2 and to assess the accuracy of this technique. Additionally, this study warrants further investigation into the effect different mutations have on binding affinity and if this is correlated with antibody escape, as this is seen in IAV (Hensley et al., 2009). To study this, SARS-CoV-2 could then be passaged in cells from either vaccinated or non-vaccinated mice and the ability of any mutant to evade antibodies and bind to ACE2 could be assessed using ELISA.

## Appendix

**Figure 4.**
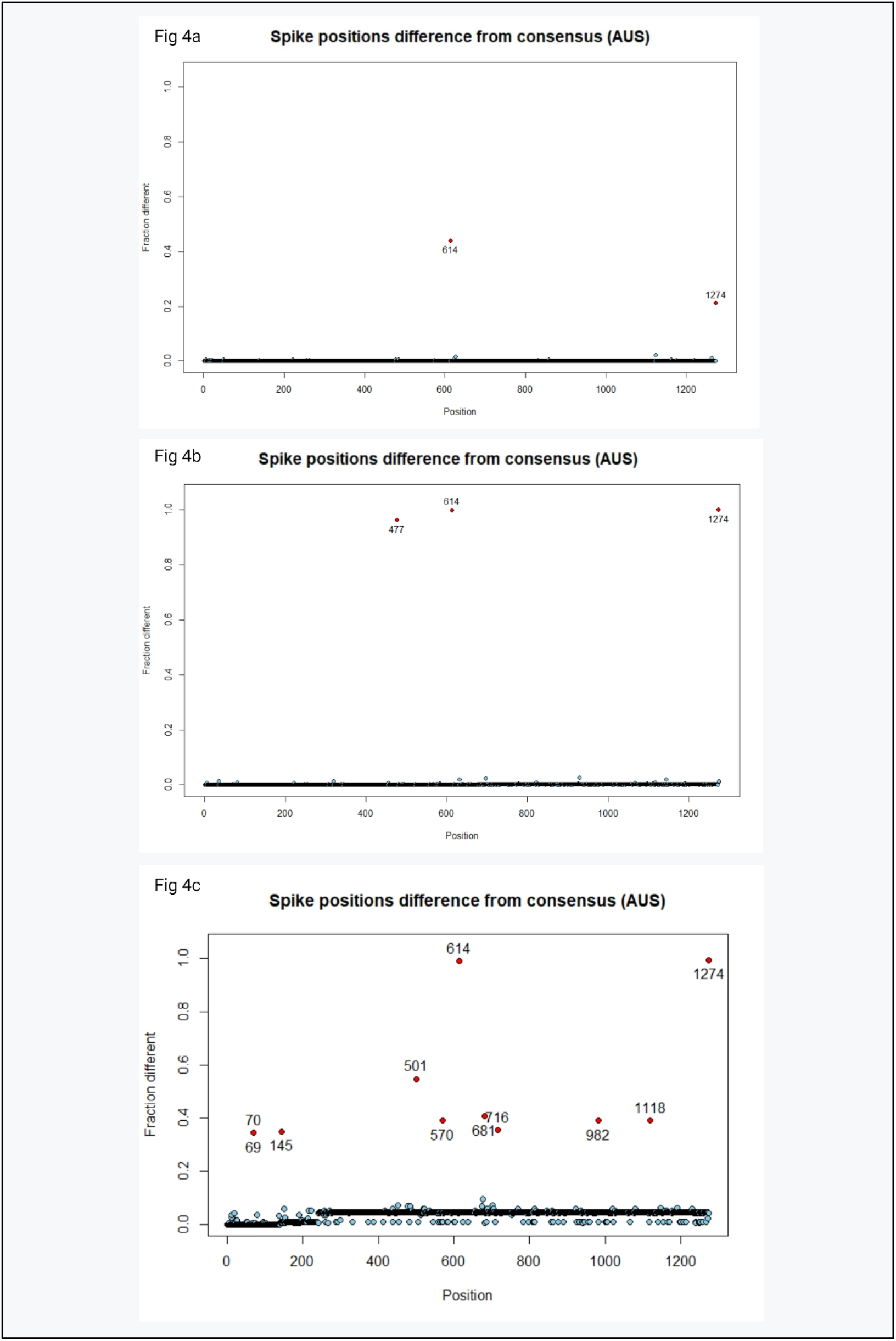
Amino acid positions which differ from the Wuhan reference sequence in sequences from Australia. Figure 4a, 4b and 4c, depict the sequences from wave 1, 2 and 3 respectively and amino acid positions in which the composition differs from the Wuhan reference sequence in more than 10% of sequences sampled are highlighted in red and labelled.

**Figure 5.**
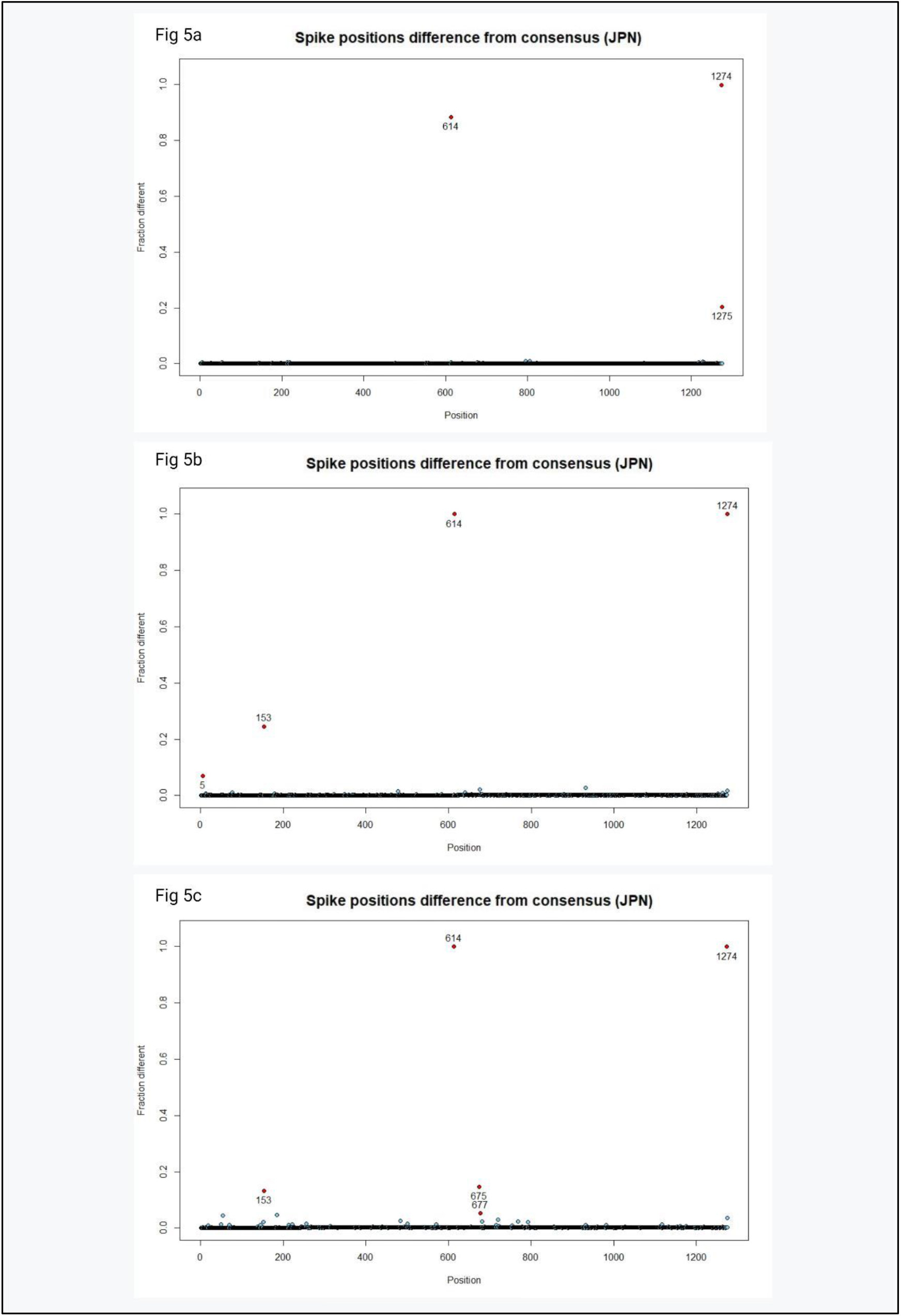
Amino acid positions which differ from the Wuhan reference sequence in sequences from Japan. Figure 5a, 5b and 5c, depict the sequences from wave 1, 2 and 3 respectively and amino acid positions in which the composition differs from the Wuhan reference sequence in more than 10% of sequences sampled are highlighted in red and labelled.

**Figure 6.**
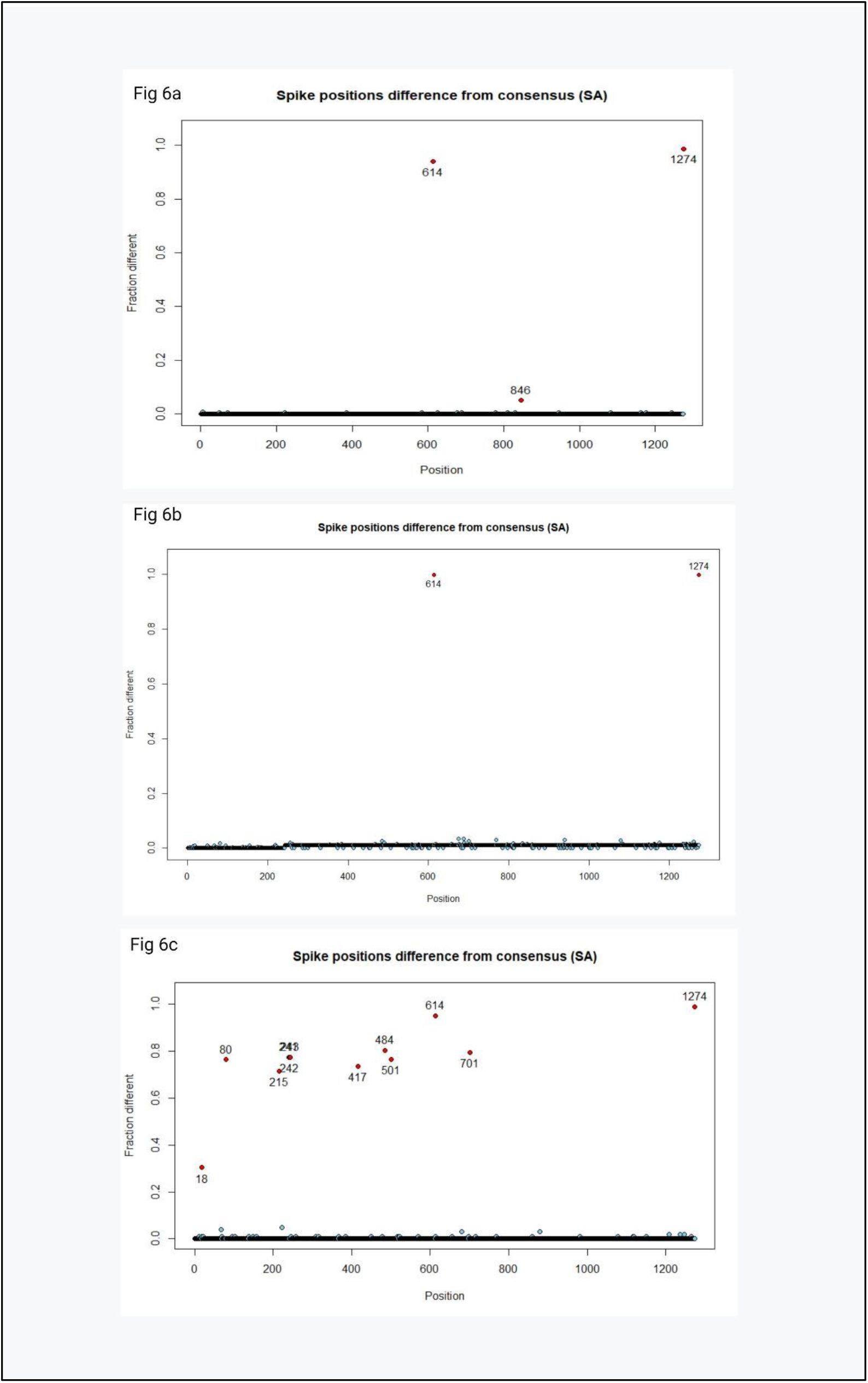
Amino acid positions which differ from the Wuhan reference sequence in sequences from South Africa. Figure 6a, 6b and 6c, depict the sequences from wave 1, 2 and 3 respectively and amino acid positions in which the composition differs from the Wuhan reference sequence in more than 10% of sequences sampled are highlighted in red and labelled.

**Figure 7.**
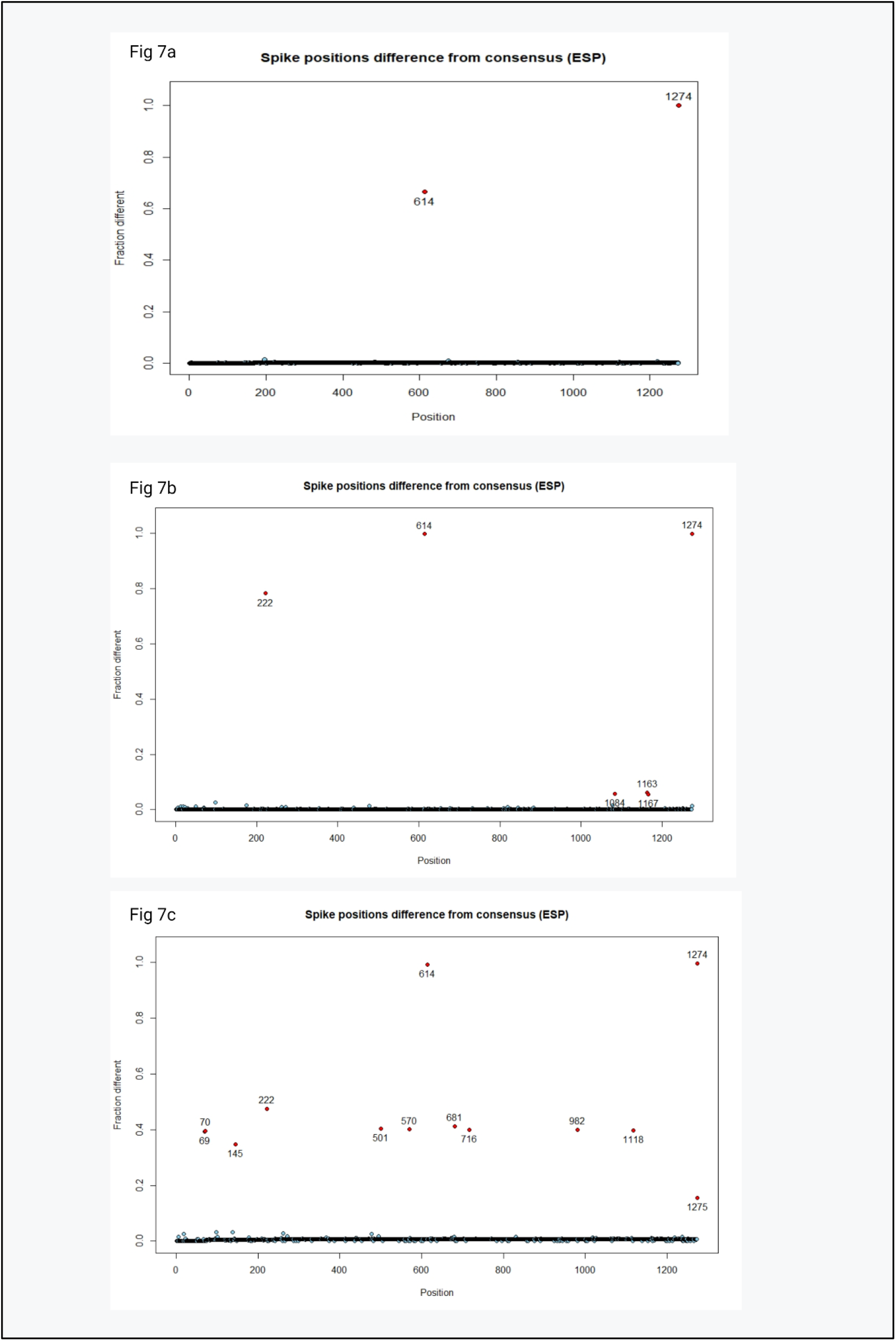
Amino acid positions which differ from the Wuhan reference sequence in sequences from Spain. Figure 7a, 7b and 7c, depict the sequences from wave 1, 2 and 3 respectively and amino acid positions in which the composition differs from the Wuhan reference sequence in more than 10% of sequences sampled are highlighted in red and labelled.

**Figure 8.**
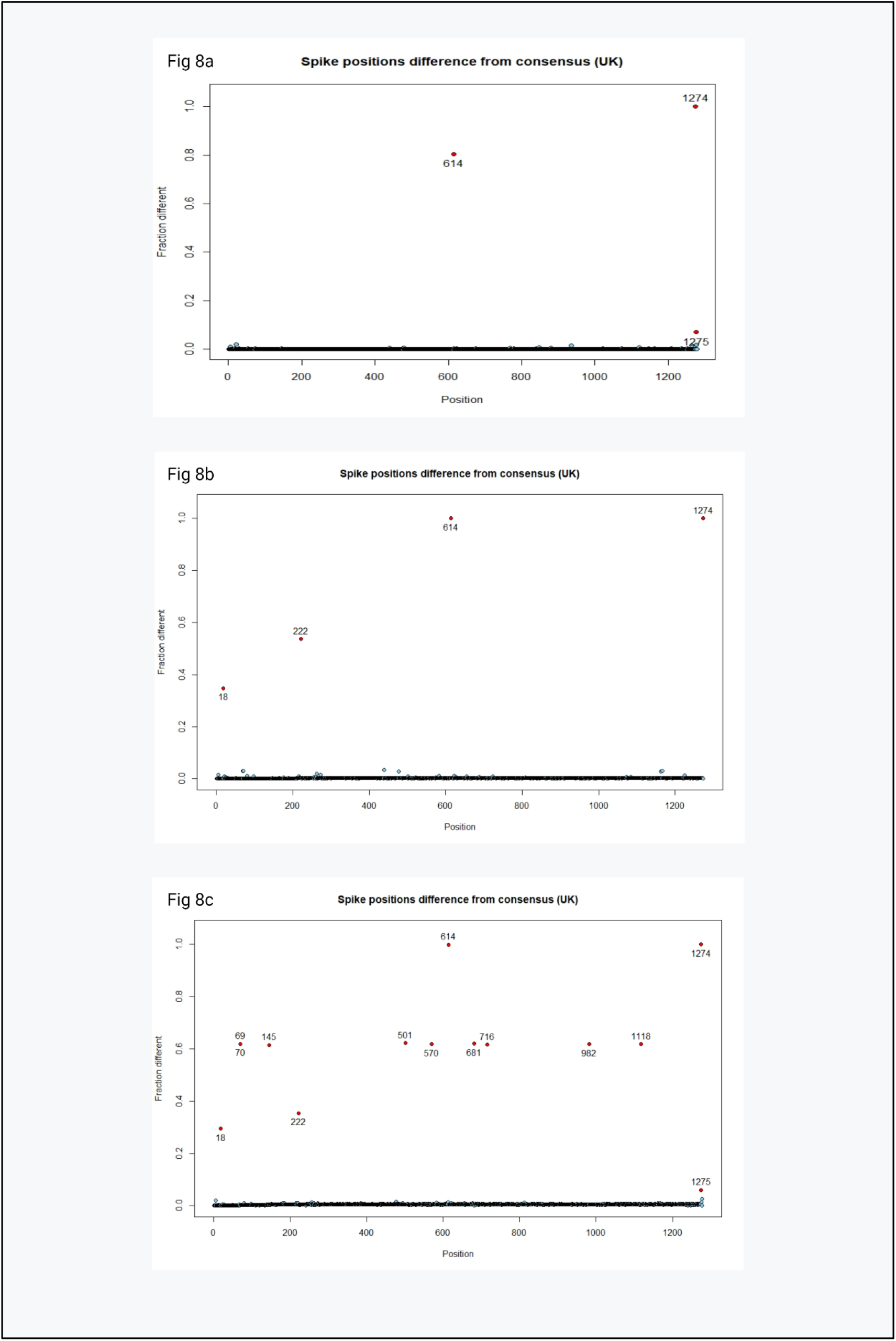
Amino acid positions which differ from the Wuhan reference sequence in sequences from the United Kingdom. Figure 8a, 8b and 8c, depict the sequences from wave 1, 2 and 3 respectively and amino acid positions in which the composition differs from the Wuhan reference sequence in more than 10% of sequences sampled are highlighted in red and labelled.

**Figure 9.**
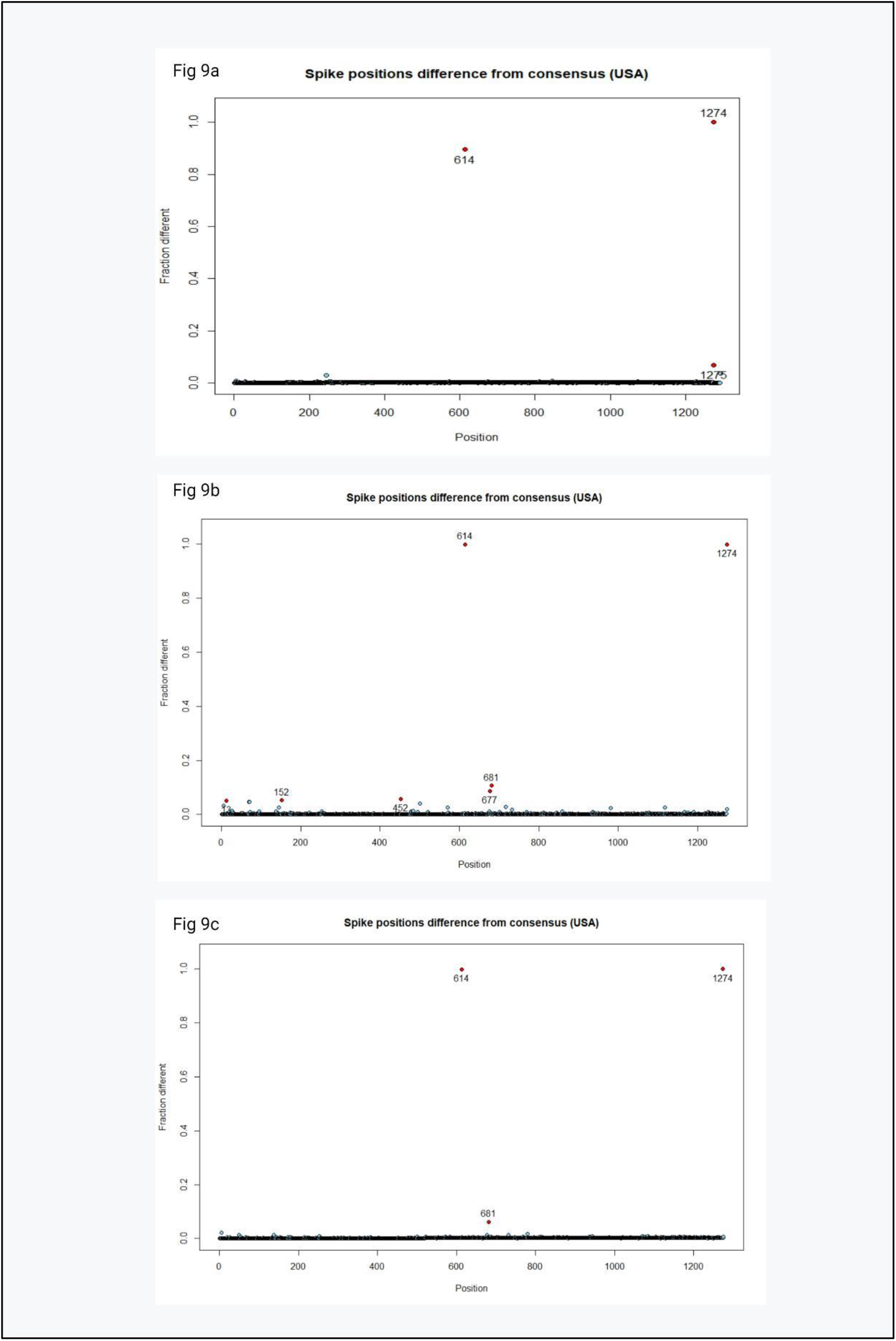
Amino acid positions which differ from the Wuhan reference sequence in sequences from the USA. Figure 9a, 9b and 9c, depict the sequences from wave 1, 2 and 3 respectively and amino acid positions in which the composition differs from the Wuhan reference sequence in more than 10% of sequences sampled are highlighted in red and labelled.

